# O-GlcNAcAtlas: A Database of Experimentally Identified O-GlcNAc Sites and Proteins

**DOI:** 10.1101/2020.11.25.397042

**Authors:** Junfeng Ma, Yaoxiang Li, Chunyan Hou, Ci Wu

**Affiliations:** Department of Oncology, Lombardi Comprehensive Cancer Center, Georgetown University Medical Center, Washington DC, USA; Dalian Institute of Chemical Physics, Chinese Academy of Sciences, Dalian, Liaoning, China

**Keywords:** *database*, O-GlcNAc, proteomics

## Abstract

O-linked β-N-acetylglucosamine (O-GlcNAc) is a post-translational modification (i.e., O-GlcNAcylation) on serine/threonine residues of proteins. As a unique intracellular monosaccharide modification, protein O-GlcNAcylation plays important roles in almost all biochemical processes examined. Aberrant O-GlcNAcylation underlies the etiologies of a number of chronic diseases (including cancer, diabetes, and neurodegenerative disease). With the tremendous improvement of techniques, thousands of proteins along with their O-GlcNAc sites have been reported. However, until now there is no database dedicated to accommodate the rapid accumulation of such information. Thus, O-GlcNAcAtlas is created to integrate all experimentally identified O-GlcNAc sites and proteins from 1984 to Dec, 2019. O-GlcNAcAtlas consists of two datasets (Dataset-I and Dataset-II, for unambiguously identified sites and ambiguously identified sites, respectively), representing a total number of 4571 O-GlcNAc modified proteins. For each protein, comprehensive information (including gene name, organism, modification sites, site mapping methods and literature references) is provided. To solve the heterogeneity among the data collected from different sources, the sequence identity of these reported O-GlcNAc peptides are mapped to the UniProtKB protein entries. To our knowledge, O-GlcNAcAtlas is the comprehensive and curated database encapsulating all O-GlcNAc sites and proteins identified in the past 35 years. We expect that O-GlcNAcAtlas will be a useful resource which will facilitate site-specific O-GlcNAc functional studies and computational analyses of protein O-GlcNAcylation. The public version of the web interface to the O-GlcNAcAtlas can be found at https://oglcnac.org.

## 1. Introduction

O-linked β-N-acetylglucosamine (O-GlcNAc), which was discovered in early 1980s, is a post-translational modification (i.e., O-GlcNAcylation) on serine/threonine residues of proteins (Torres et al. 1984; Holt et al. 1986). Distinct from the traditional glycosylation (i.e., N-glycosylation, O-glycosylation, and GPI-anchored glycosylation), O-GlcNAcylation is a unique intracellular monosaccharide modification without being further elongated into complex sugar structures (Wells et al. 2001; Hart et al. 2007). By modulating various aspects of target proteins (e.g., activity, localization, stability and others), O-GlcNAcylation exerts diverse functional roles (Hart et al. 2011; Bond et al. 2013; Hart 2019). After several decades’ endeavor, it has been revealed that O-GlcNAcylation exists in all metazoans (including animals, insects, and plants), some bacteria, fungi and virus. For example, mounting evidence has demonstrated that deregulated protein O-GlcNAcylation underlies multiple human diseases, especially in diabetes (Ma et al., 2013; Vaidyanathan et al. 2014), cancer (Slawson et al. 2011; Ma et al. 2014; Ferrer et al. 2016), and neurodegenerative diseases (Yuzwa et al. 2014; Wani et al. 2017). Moreover, targeting protein O-GlcNAcylation holds great promise for biomedical applications (e.g., as therapeutic targets and biomarkers) (Zhu et al. 2020).

Although great progress has been made towards the understanding of myriads roles of protein O-GlcNAcylation, site-specific O-GlcNAc studies have been lagged behind, largely due to lack of powerful site mapping methods. Indeed, low throughout methods (e.g., Edman degradation and site-directed mutagenesis) played pivotal roles for O-GlcNAc identification on proteins of interest in the early days. With the development of enrichment and identification techniques in recent years, mass spectrometry-based proteomics began to be exploited as a sensitive and high throughput tool for large-scale identification of O-GlcNAc proteins (Wang et al. 2008; Ma et al. 2014; Thompson et al. 2018). It becomes possible to identify tens of hundreds of O-GlcNAc sites in one single experiment by using proteomics (Wang et al. 2010; Zhao et al. 2011; Trinidad et al. 2012; Alfaro et al. 2012; Ma et al. 2015; Wang et al. 2017; Xu et al., 2017; Woo et al. 2018; Qin et al. 2018; Li et al. 2019).

Although there are a number of databases developed for other PTMs, including dbPTM (Huang et a;. 2019), O-GlycBase (Gupta et al. 1999), UniCarbKB (Campbell et al. 2014), GlyGen (York et al. 2020), Glycosciences.DB (Bohm et al. 2019), GlyTouCan (Tiemeyer et al. 2017), and PhosphoSite Plus (Hornbeck et al. 2019). Until now a few databases have been created to specifically accommodate the rapid accumulation of O-GlcNAc information on proteins. The database of O-GlcNAcylated proteins and sites (dbOGAP) which was constructed in 2011 contains ~400 O-GlcNAcylation sites and has not been updated (Wang et al. 2011). Undoubtedly, there is an urgent need to create a comprehensive and curated O-GlcNAc-specific database. Herein, we describe O-GlcNAcAtlas, a manually curated database of experimentally identified O-GlcNAc sites and proteins in the past decades. By enabling users to search and retrieve data easily, O-GlcNAcAtlas is proposed to facilitate site-specific functional analysis of O-GlcNAc proteins.

## 2. Methods

The system flow of the construction of the O-GlcNAcAtlas is presented in Figure 1. Specifically, O-GlcNAcAtlas was compiled through a manual curation of the literature accessed from PubMed between 1984 and Dec. 30, 2019. The following search items: ‘O-linked β-N-acetylglucosamine’, ‘O-GlcNAc’, or ‘O-GlcNAcylation’ were used. O-GlcNAc sites information in each publication was retrieved and evaluated by at least one of the curators. Besides O-GlcNAc sites, related information (including species, sample type, peptide sequence, protein name and site-mapping methods used) was also extracted. To determine the positions of O-GlcNAcylated Ser/Thr residues, the experimentally identified peptides were then mapped to UniProtKB protein entries based on database identifier or sequence similarity. The O-GlcNAcylated peptides/sites that could not align exactly to a protein sequence were annotated with curators’ comments.

**Figure 1.**
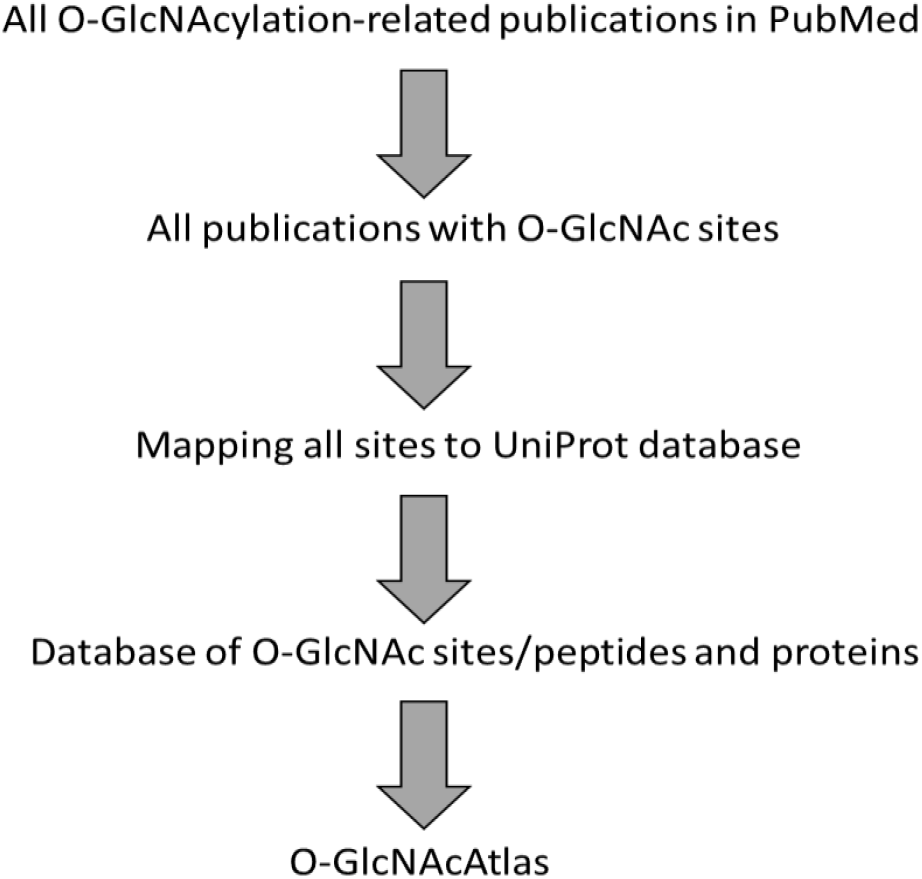
Assembly of experimentally identified O-GlcNAc sites and proteins for a comprehensive database O-GlcNAcAtlas.

Finally, each mapped O-GlcNAc site was attributed to the corresponding literature (PubMed ID). Of special note, to avoid and minimize misleading and confusion, rigorous selection criteria was applied to the O-GlcNAc sites and proteins selected. For large-scale proteomics studies, proteins without O-GlcNAc peptides/sites identified were not included. Each entry from low-throughput studies were also carefully curated.

A user-friendly web-based graphical user interface was created with HTML, CSS, and Bootstrap. The backend server is running on a collection of services developed using Python programming language (version 3.8.1) and coupled with the MySQL database. All entries, given a unique O-GlcNAcAtlas identification number, were organized in the MySQL database.

## 3. Current state of O-GlcNAcAtlas

Literature mining from PubMed yielded a total of 2236 O-GlcNAc-relevant articles (Figure 2A). Among them, 225 articles contain O-GlcNAc sites on proteins (Figure 2A). Each publication was retrieved and evaluated by at least one of the curators, with O-GlcNAc sites and related recorded and compiled. Clearly there has been increased interest in O-GlcNAc studies, especially site-specific functional studies in the past decade.

**Figure 2.**
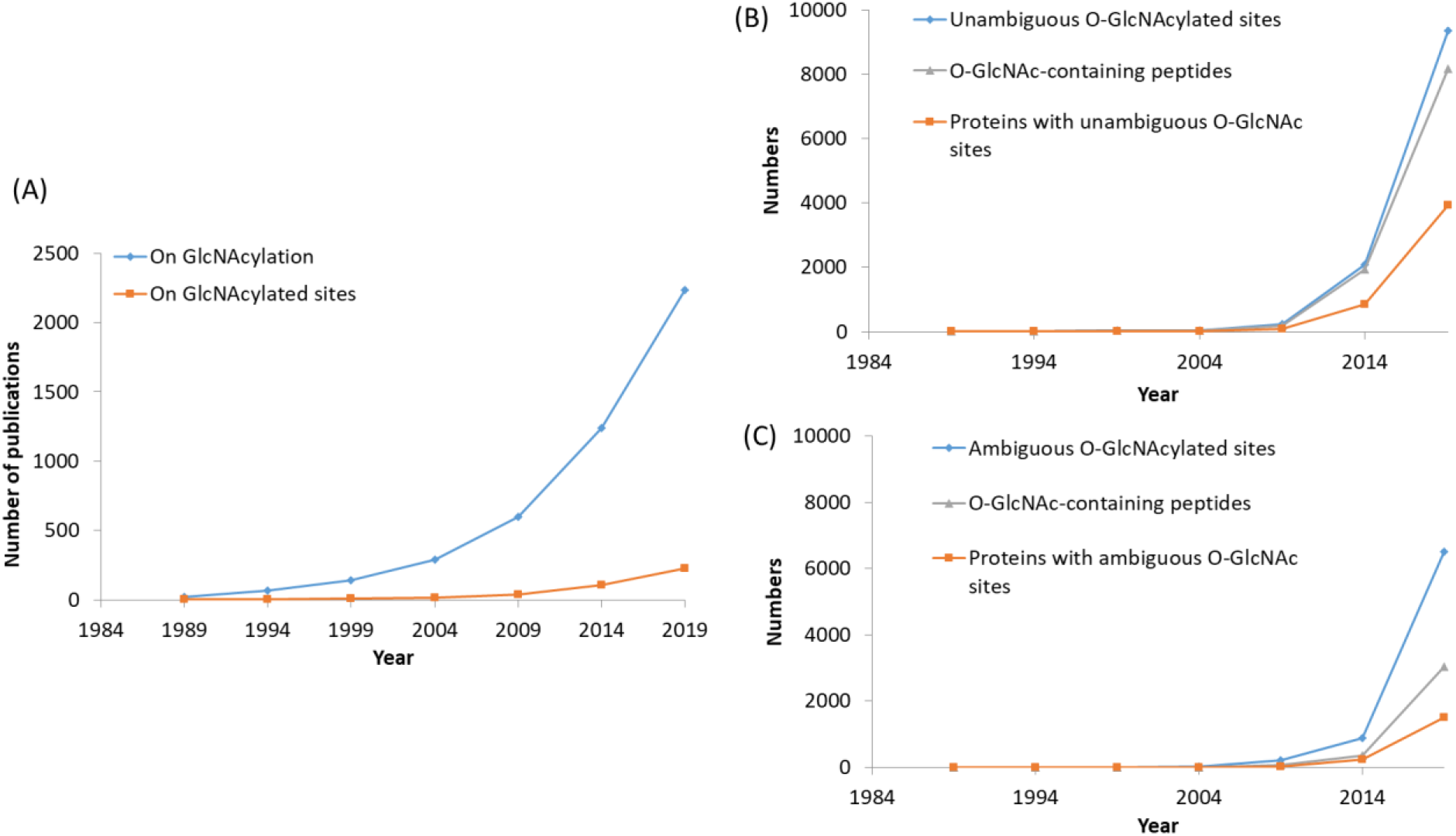
(A) The accumulated number of O-GlcNAcylation-related publications and publications identifying O-GlcNAc sites from 1984 through Dec. 2019. Accumulation of unambiguous O-GlcNAc sites (B) and ambiguous O-GlcNAc sites (C) as well as their corresponding peptides and proteins identified from 1984 through Dec. 2019.

O-GlcNAcAtlas consists of two datasets, depending on the ambiguity of O-GlcNAc sites mapped. Dataset-I contains unambiguously assigned O-GlcNAc sites, while Dataset-II is for O-GlcNAc sites ambiguously identified (mainly due to the low localization scores by software tools especially for peptides with clustered Ser/Thr residues). Despite the ambiguity of specific modification sites, the corresponding peptides can be positively identified, so do the O-GlcNAc proteins. Considering Dataset-2 provides useful information, it has been kept into the database. Overall, 9348 O-GlcNAc sites were unambiguously identified, corresponding to 8151 peptides and 3918 proteins (Figure 2B). In addition, 3028 peptides on 1507 proteins were found to be O-GlcNAcylated, corresponding to ~6520 ambiguous sites (Figure 2C).

Among the 9348 unambiguous O-GlcNAc sites, >98% were identified during 2010-2019 (Figure 3A). Moreover, >98% of all sites were unambiguously assigned by mass spectrometry (Figure 3B). While ~15% of all sites were identified by two or more publications, the majority (85%) were found only once (Figure 3C). And it turns out that the distribution of Ser residues and Thr residues is 62%:38% (slightly less than a ratio of 2:1) (Figure 3D).

**Figure 3.**
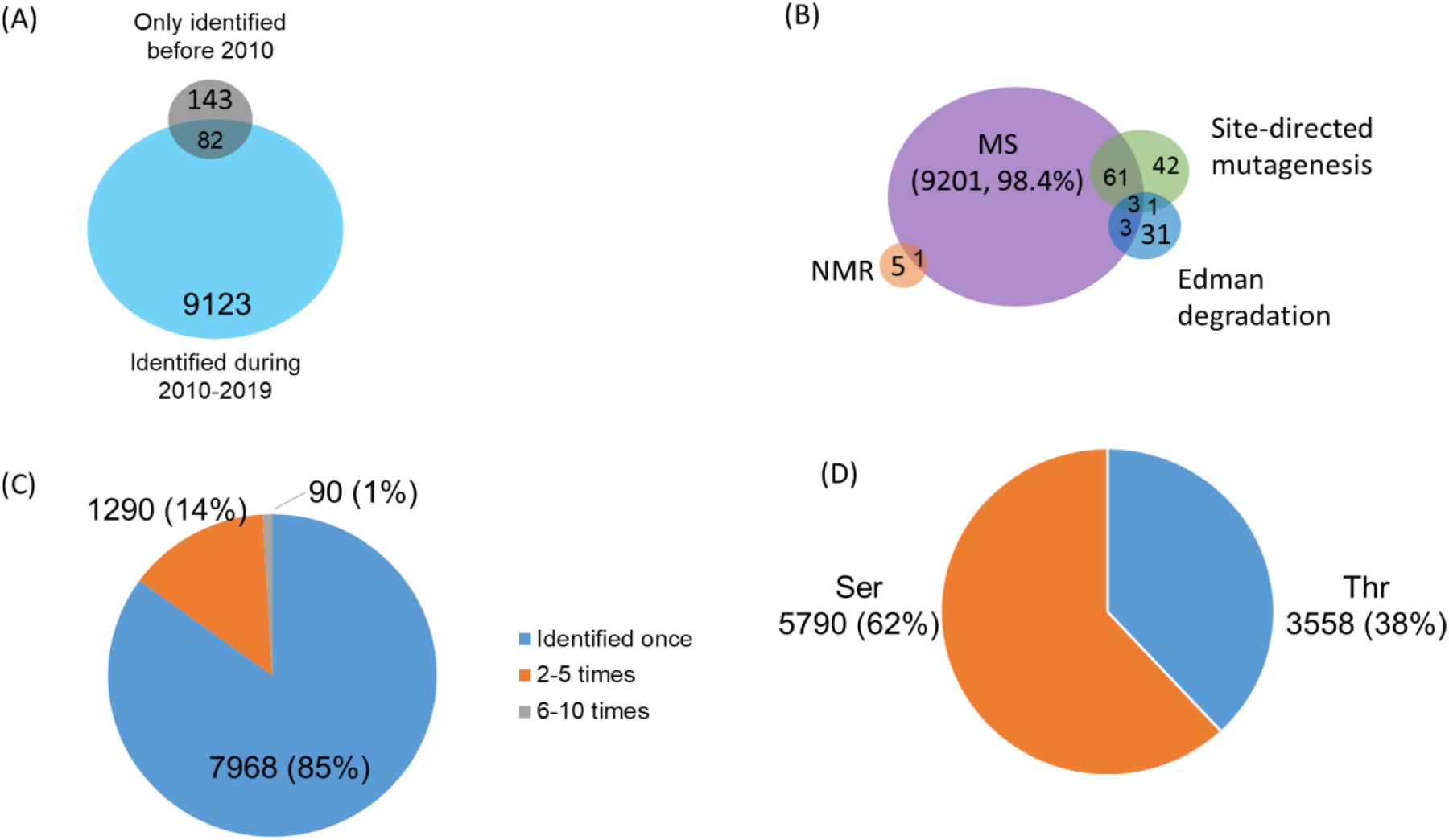
Distribution of unambiguously identified O-GlcNAc sites. (A) Classification of O-GlcNAc sites according to their year of publication. (B) O-GlcNAc sites by different identification methods. (C) The identification frequencies of the O-GlcNAc sites by mass spectrometry. (D) Distribution of Ser/Thr residues modified by O-GlcNAc. (Note: MS, mass spectrometry; NMR, nuclear magnetic resonance spectroscopy)

Besides 3918 proteins with unambiguous O-GlcNAc sites, 1507 proteins were matched with ambiguous O-GlcNAc sites. Providing 854 proteins were overlapped between the two sets, in total 4571 O-GlcNAc proteins were identified (Figure 4A). Among the O-GlcNAc proteins, ~77% (3535 out of 4571 proteins) were identified by one study (Figure 4B). However, 27 proteins were identified at least 10 times (Table 1). Although ~62% of proteins are derived from human, 38% (1728 out of 4571 proteins) are from other organisms (mainly common model systems, such as mouse, rat, C. elegans, Drosophila, Arabidopsis, and wheat) (Figure 4C). The details of O-GlcNAc proteins/sites information from different species are shown in Table 2. Regarding human proteins, most O-GlcNAc proteins were identified from model cell lines (e.g., HeLa cells and HEK293 cells) (Figure 4D). Moreover, hundreds of O-GlcNAc proteins were also identified from tissues/cells of special research interest (e.g., primary T cells and brain).

**Figure 4.**
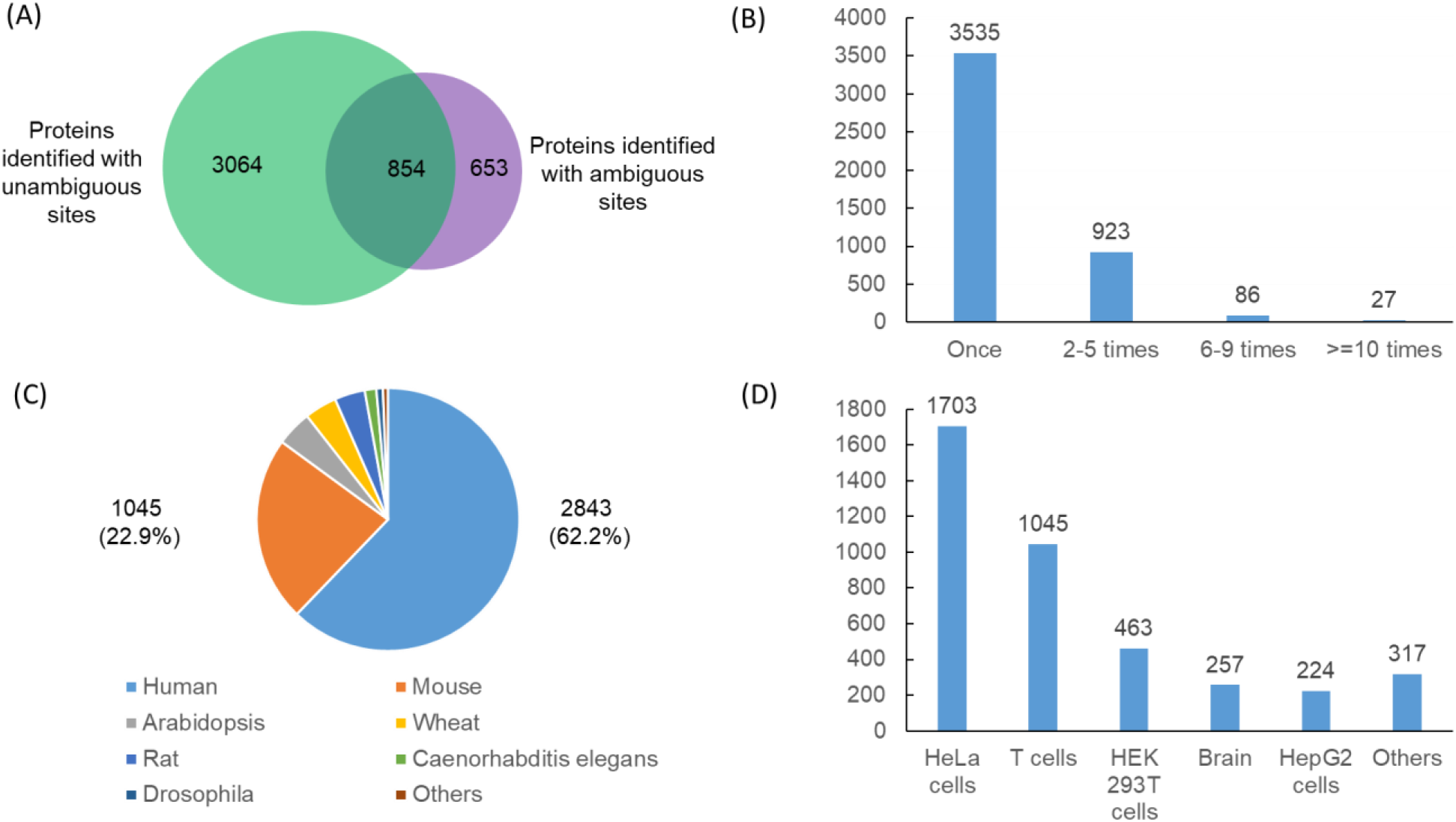
Distribution of 4571 O-GlcNAcylated proteins. (A) Proteins matched with unambiguous sites and ambiguous sites. (B) A representation of the number of times a specific protein is identified. The majority of proteins (77%) are only identified by one publication. However, 27 proteins were identified at least 10 times (listed in Table 1). (C) Distribution of proteins in different species. (D) Number of human proteins identified from human cultured cells and other sources studied.

**Table 1.**
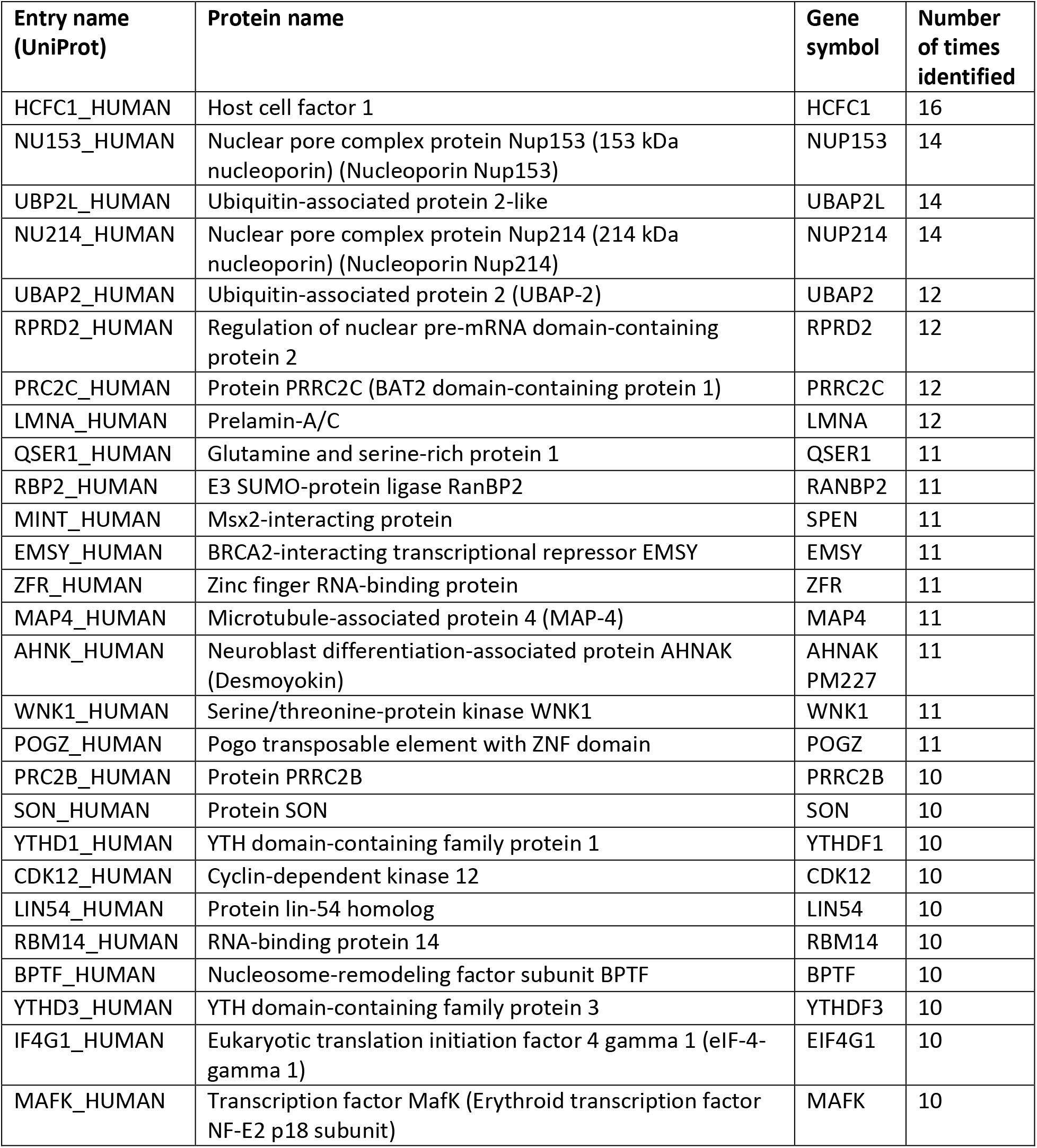
A list of 27 proteins identified independently in at least 10 publications.

**Table 2.** Summary of O-GlcNAc sites and proteins identified from different species.

| Species | Dataset-I |  | Dataset-II |  | Total proteins |
| --- | --- | --- | --- | --- | --- |
|  | Unambiguous sites | Proteins matched with unambiguous sites | Ambiguous Sites | Proteins matched with ambiguous sites |  |
| Human | 5654 | 2273 | 5074 | 1202 | 2843 |
| Mouse | 2315 | 1017 | 573 | 98 | 1045 |
| Arabidopsis | 334 | 167 | 559 | 138 | 200 |
| Wheat | 386 | 182 |  |  | 182 |
| Rat | 428 | 159 | 84 | 21 | 171 |
| Caenorhabditis elegans | 66 | 57 | 88 | 11 | 65 |
| Drosophila | 103 | 36 | 131 | 33 | 37 |
| Others | 62 | 27 | 11 | 4 | 28 |

## 4. O-GlcNAcAtlas web-server

To facilitate the use of the O-GlcNAcAtlas resource, a web interface has been developed for users to browse and search efficiently for their O-GlcNAcylated proteins of interest. O-GlcNAcAtlas can be searched using UniProt accession, protein name, or gene symbol as key words, and the results can be filtered further. The search output includes the basic annotations for all the matched entries (Figure 5A). The accession number of each entry is linked to the detailed annotation for the specific protein (Figure 5B). So far, O-GlcNAcAtlas supports several functions including data searching, browsing and retrieving. Moreover, search results can be directly downloaded and saved from the O-GlcNAcAtlas webpage.

**Figure 5.**
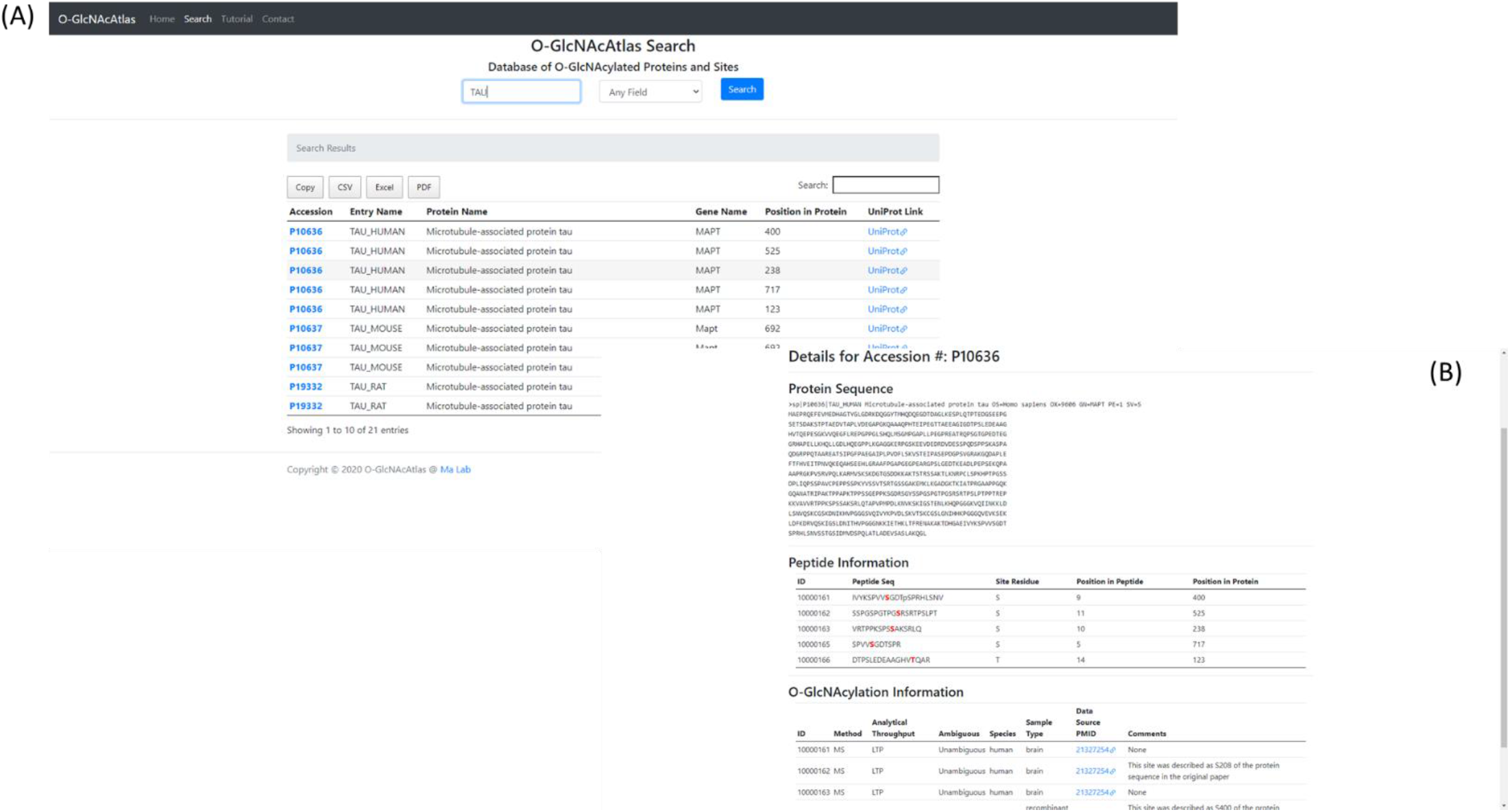
A snapshot for searching O-GlcNAcAtlas, with ‘microtubule-associated protein tau’ as an example. (A) Tabular results for all the matched entries. (B) Main display page with detailed annotation and links to UniProtKB and PubMed.

## 5. Conclusion

To appreciate the tremendous efforts in O-GlcNAc research in the past 35 years, we aimed to create a comprehensive database of O-GlcNAc sites and proteins. O-GlcNAcAtlas not only includes data from case-by-case studies but also integrates high-throughput data from proteomics studies. For either low-throughput or high-throughput studies, we tried our best to carefully curate each entry, with curators’ comments added. We fully respect the original authors’ discoveries, but by adding curators’ comments we hope viewers can pay attention to specific entries (e.g., the erroneously labeled modification residues or sites). With O-GlcNAcAtlas, we aim to provide a one-stop portal for the community to search O-GlcNAcylated proteins and the sites information. We anticipate it will facilitate both basic and translational research to better understand protein O-GlcNAcylation at the molecular level.

## Acknowledgements

We are indebted to Dr. Gerald Hart for his insights during the initiation of this project several years ago and his continuous support along the years. We wish to thank Dr. Zhangzhi Hu (at NIH) and Dr. Leslie Arminski and Dr. Hongzhan Huang (at Protein Information Resource) for helpful discussions at the beginning of this project. We appreciate the kind encouragement and comments from Dr. Michelle Bond and many other peers. We would like to acknowledge researchers who, by generously answering our curators’ questions regarding O-GlcNAc sites in specific publications, have contributed to improve the database. Last but not least, we would appreciate if investigators can report any potentially missing sites and/or send the datasets in their new publications to us so that we can update this database in a timely manner to further facilitate researchers in the O-GlcNAc field.

## Abbreviation

O-GlcNAc: O-linked β-N-acetylglucosamine
MS: mass spectrometry
NMR: nuclear magnetic resonance spectroscopy

